# Monocytes mediate *Salmonella* Typhimurium-induced tumour growth inhibition

**DOI:** 10.1101/2020.05.13.092858

**Authors:** Síle A. Johnson, Michael J. Ormsby, Hannah M. Wessel, Heather E. Hulme, Alberto Bravo-Blas, Anne McIntosh, Susan Mason, Seth B. Coffelt, Stephen W.G. Tait, Allan McI. Mowat, Simon W.F. Milling, Karen Blyth, Daniel M. Wall

**Affiliations:** Institute of Infection, Immunity and Inflammation, College of Medical, Veterinary and Life Sciences, Sir Graeme Davies Building, University of Glasgow, Glasgow G12 8TA, United Kingdom; Cancer Research UK Beatson Institute, Garscube Estate, Switchback Road, Glasgow G61 1BD, United Kingdom; Institute of Cancer Sciences, University of Glasgow, Bearsden Road, Glasgow, G61 1QH, United Kingdom

**Keywords:** Monocytes, SL7207, bacterial cancer therapy, immunotherapy

## Abstract

The use of bacteria as an alternative cancer therapy has been re-investigated in recent years. A number of bacterial strains for this purpose have been generated, one of which is SL7207: an auxotrophic *Salmonella enterica* serovar Typhimurium *aroA* mutant with immune-stimulatory potential. Here we show that systemic administration of SL7207 induces melanoma tumour growth arrest *in vivo*, with greater survival of the SL7207-treated group compared to control PBS-treated mice. Administration of SL7207 is accompanied by a change in the immune phenotype of the tumour-infiltrating cells towards pro-inflammatory, with expression of the T_H_1 cytokines IFN-γ, TNF-α, and IL-12 significantly increased. Interestingly, Ly6C^+^MHCII^+^ monocytes were recruited to the tumours following SL7207 treatment and were pro-inflammatory. Accordingly, the abrogation of these infiltrating monocytes using clodronate liposomes prevented SL7207-induced tumour growth inhibition. These data demonstrate a previously unappreciated role for infiltrating inflammatory monocytes underlying bacterial-mediated tumour growth inhibition. This information highlights a novel role for monocytes in controlling tumour growth, contributing to our understanding of the immune responses required for successful immunotherapy of cancer.

## Introduction

Since the late 19^th^ century, bacteria and bacterial products have been studied and used clinically to treat solid tumours (Coley, 1893). There is much evidence to support the anti-tumour effects of bacteria such as *Salmonella enterica* serovar Typhimurium, *Escherichia coli, Listeria monocytogenes, BCG* and others in *in vivo* tumour models (Crull *et al*., 2011; Lizotte *et al*., 2014; Morales *et al*., 1976; Stern *et al*., 2015; Zheng *et al*., 2017). These bacteria can selectively accumulate in the tumour microenvironment and induce anti-tumour effects through a number of mechanisms such as metabolite depletion and direct cell killing (Chen *et al*., 2012; Fu *et al*., 2008; Kasinskas & Forbes, 2006; Kasinskas & Forbes, 2007). A primary mechanism which has been the subject of intense study is immune infiltration into the tumour following bacterial administration into tumour-bearing mice, with many immune cells implicated in the anti-tumour effects which are seen to occur. Dendritic cells are recruited to infected tumours following *S*. Typhimurium administration, and those isolated from the tumour-draining lymph nodes produced more IL-6, TNF-α and IL-1β than dendritic cells harvested from control mice (Avogadri *et al*., 2005, 2008). Furthermore, *S*. Typhimurium treatment of tumour-bearing mice has resulted in the increased expression of connexin 43, a gap junction pore-forming protein which facilitates the movement of tumour-associated antigen from tumour cells to dendritic cells (Saccheri *et al*., 2010). Lymphocyte involvement following bacterial administration to tumour-bearing mice has revealed somewhat contradictory findings. In one study, the depletion of CD4 using anti-CD4 antibodies did not significantly affect the tumour growth inhibitory effects of *Escherichia coli*, whilst the blockage of CD8 completely abolished anti-tumour efficacy (Stern *et al*., 2015). However, previous work indicated a depletion of either CD4 or CD8 alone mildly abrogated the anti-tumour effects of *Salmonella enterica* serovar *Choleraesuis* (Lee *et al*., 2011). Furthermore, given these reports, and others which have claimed a role for T cells in mediating the anti-tumour effects for bacteria, it is somewhat surprising that bacteria are also capable of tumour growth inhibition in athymic nude mice (Zhang *et al*., 2015; Zhao *et al*., 2005; Zhao *et al*., 2006).

Monocytes and macrophages have garnered little attention in the bacterial-meditated cancer therapy literature. This is interesting given the critical role the cells play in mediating the immune response to oral *S*. Typhimurium infection (Rydström & Wick, 2007, 2009; Yrlid *et al*., 2000). These immune responses include a host of anti-tumour immune cells being recruited and producing an array of cytokines such as TNF-α and IL-12, which are not conducive to established tumour growth (Boggio *et al*., 2000; Brunda *et al*., 1993; Sabel *et al*., 2004). However, it has been reported that there was an increase in the number, or density, of macrophages in the tumours following systemic bacterial administration (Lee *et al*., 2011; Lizotte *et al*., 2014). For one of these studies, the only marker used to identify macrophages was CD11b, which would also include other myeloid cells such as dendritic cells and neutrophils, thus making conclusions about the role of macrophages in this setting difficult (Lizotte *et al*., 2014). More recently, F4/80^+^ cells were credited with a pro-inflammatory role in the tumour following infection with *S*. Typhimurium (Zheng *et al*., 2017). However, as F4/80 stains both macrophages and monocytes transitioning to macrophages, it is again difficult to determine the role the individual cell types play in the inflammatory signature and associated tumour regression.

Monocytes have yet to be thoroughly investigated for their anti-tumour effects in the context of cancer immunotherapy strategies. It is credible that monocytes should accumulate in the infected tumour as these cells have been shown to accumulate in inflamed tissues and provide inflammatory cytokines such as TNF-α which would not be conducive to tumour progression (Bain *et al*., 2013; Kocijancic *et al*., 2017; Sabel *et al*., 2004; Zigmond *et al*., 2012). The present study sought to address this dearth of information pertaining to tumour-infiltrating monocytes following systemic administration of SL7207, an attenuated strain of *S*. Typhimurium (Hoiseth & Stocker, 1981; Johnson *et al*., 2017). Here we demonstrate that these monocytes accumulate in the tumour following systemic infection, are highly pro-inflammatory and are required for the anti-tumour effects of SL7207, highlighting an important role for these cells in anti-tumour immune responses.

## Methods

### Bacterial strains

*Salmonella enterica* serovar Typhimurium SL7207 (Δ*aroA*) was kindly provided by Prof Siegfried Weiss (Helmholtz Centre for Infection Research) (Hoiseth & Stocker, 1981; Johnson *et al*., 2017). Bacterial cultures were maintained on Lysogeny broth (LB) agar supplemented with appropriate antibiotics. Luciferase expressing (*lux*) SL7207 (SL-*lux*) was generated using the method of (Riedel *et al*., 2007).

### Cell lines and animals

The B16F10 mouse melanoma cell line was kindly provided by Prof Gerry Graham (University of Glasgow) and cells were maintained in Dulbecco’s Modified Eagle Serum (DMEM; Gibco^®^, 12491) supplemented with 10% fetal calf serum (FCS), 1 mM L-glutamine, 2 mM sodium pyruvate and 100 international units (I.U.)/ml penicillin/streptomycin at 37°C and 5% CO_2_. All cells were routinely tested for mycoplasma contamination. Five to eight-week-old female C57BL/6 mice were purchased from Charles River Laboratories. All animal procedures were approved by internal University of Glasgow and Beatson Institute ethics committee and were carried out in accordance with the relevant guidelines and regulations as outlined by the UK Home Office (PPL70/8584 and PPL70/8645).

### Infection of tumour-bearing mice and recovery of bacteria from tissues

Six to nine-week-old female C57BL/6 mice were inoculated subcutaneously in the right back flank with 2 x 10^5^ B16F10 cells in cold, sterile phosphate buffered saline (PBS). Mice bearing melanoma tumours of greater than 200 μm^3^ (8-9 days post tumour cell transfer) were intravenously (i.v.) injected with 5 x 10^6^ colony forming units (CFU) SL7207-*lux* in 100 μl or 100 μl sterile PBS as a control. Tumour growth was measured throughout with Vernier callipers. For the recovery of bacteria, melanoma tumour, spleen and liver were carefully resected, weighed and placed in ice cold PBS. Tissues were then placed in 1-2 ml of ice-cold PBS in 5 ml bijoux and homogenised using a hand held tissue homogeniser (OMNI International Inc., TM125-220). Homogenates were then serially diluted in PBS and these dilutions were plated out on LB agar. Plates were checked for bioluminescent light emission to ensure SL7207-*lux* isolation using the *in vivo* imaging system (IVIS; Perkin Elmer).

### Flow cytometry

Tissues were carefully resected, weighed, and placed in ice cold PBS. Tumours were then transferred to digestion medium composed of 3 mg/ml collagenase A (Sigma, 10103586001) and 25 μg/ml DNAse I (Sigma, 10104159001) in DMEM (Coffelt *et al*., 2015). All tissue was digested at 37°C for 30 min with intermittent vigorous shaking before being passed through a 70 μm strainer (VWR, 734-0003) and neutralized with 8% FCS-DMEM buffer. Tumour cells were treated with 1 ml red blood cell lysis buffer (Sigma, 11814389001) for 5 min at room temperature, neutralised with 10 ml of 8% FCS-DMEM and were centrifuged at 400 x *g* for 5 minutes at 4°C. Cells were resuspended in flow cytometry buffer (FB: 2% FCS, 3 nM EDTA (Sigma, E9884) in PBS). A portion of the cells were counted using Trypan Blue exclusion dye (Sigma, T8154). Single cell suspensions were washed and resuspended in FB. Cells were first stained in Fixable Viability Dye eFluor^®^ 780 (eBioscience) in PBS for 15-20 min on ice, in the dark. Following this incubation, cells were washed in 5 ml of FB and centrifuged at 400 x *g* for 5 min at 4°C. Cells were resuspended in their residual buffer before being incubated with anti-CD16/CD32 (‘Fc Block’) to reduce non-specific binding to Fc receptors (Biolegend). After 5 min, 100 μl of extracellular antibody mixes (final dilution of 1:200 for each) were added to the cells and left on ice for 20 minutes before being washed with PBS and pelleted as before. Antibodies used were against: CD11b (Clone M1/70, Biolegend), CD11c (clone N4/18, Biolegend), F4/80 (clone BM8, Biolegend), Ly6G (clone 1A8, Biolegend), MHCII (clone M5/114.15.2, Biolegend), CD45 (clone 30-F11, Biolegend), Ly6C (clone HK1.4, eBioscience) and SiglecF (clone E50-2440, BD Biosciences). Cells were analysed using the FACS AriaIII, LSRII analyser or Fortessa analyser (all BD Biosciences). All data generated was analysed using FlowJo software (Tree Star Inc, Oregon, USA).

### Intracellular cell stimulation and staining

Single cell suspensions were washed and resuspended in FB. For intracellular cytokine analysis, cells were resuspended in 500 μl of eBioscience™ Cell Stimulation Cocktail (plus protein transport inhibitors) in RPMI medium supplemented with 10% FCS, 1% L-glutamine and 0.01% β-mercaptoethanol. These cells remained at 37°C, 5% CO_2_ for 4 h before being washed in FB, centrifuged at 400 x *g* for 5 minutes at 4°C and subjected to surface staining as above. Following overnight incubation in Fix/Perm buffer, cells were washed twice with 2 ml Perm Buffer (both Foxp3/Transcription Factor staining buffer set, eBioscience). Cells were then resuspended in 200 μl of Perm Buffer supplemented with intracellular antibodies (1:200 for each) for 1 h in the dark. Antibodies used were against: Ki67 (clone 16A8, Biolegend), IFN-γ (clone 554413, BD Biosciences), IL-6 (clone MP520F3, BD Biosciences), TNF-α (clone MP6-XT22, Biolegend), IL-12p23 (clone C15.6, Biolegend), pro-IL-1β (clone NJTEN-3, eBioscience). Appropriate isotype controls for intracellular stains were included in all experiments.

### ELISA

For the sandwich ELISA protocol, supernatants were harvested from tumour cell suspensions which were stimulated *in vitro* with eBioscience™ Cell Stimulation Cocktail in RPMI medium supplemented with 10% FCS, 1% L-glutamine and 0.01% β-mercaptoethanol for 4 h. Supernatants were normalised to protein concentration before being subject to an ELISA protocol or TNF-α (Biolegend), IL-12 (Biolegend) or IFN-γ (Biolegend), according to the manufacturer’s instructions. Briefly, ELISA plates (Nunc™ Maxisorp™) were coated overnight with 100 μl of the relevant capture antibody at 4°C. Plates were washed three times with 0.05% PBS-Tween (PBS-T) and incubated with 200 μl of Blocking Solution for 1 h at room temperature before being washed 3 times. Samples were then added to the plates, 100 μl of each in duplicate, as well as the diluted standards. Plates were incubated for 2 h at room temperature before being washed 3 times with PBS-T. Following these washes, avidin-horseradish peroxidase conjugate in blocking buffer (1:500 dilution) was added to each well and incubated at room temperature for 30 minutes, before plates were stringently washed 5 times with PBS-T. TMB Substrate Reagent (1:1 mixture of Reagent A and Reagent B) was added to each well in 100 μl for up to 60 min for colour development. The optical density of the plates was read at 450 nm on a FLUOstar OPTIMA Microplate reader (BMG Labtech).

### Clodronate depletion

C57BL/6 mice were inoculated with B16F10 tumour cells as described above. Four days prior to infection, mice were administered 100 μl of Clodronate Liposomes (Liposoma) via tail vein injection (Gazzaniga *et al*., 2007; Griesmann *et al*., 2017; Rooijen & Sanders, 1994). This corresponds to an approximate concentration of 0.5 mg/20 g mouse weight. At 1 day prior to infection and 1, 3 and 5 days post-infection (dpi), mice received 200 μl Clod Lipo. At these time points, control PBS Lipo was also administered to control PBS Lipo mice. At 7 dpi, tissues were harvested and processed as described.

### Statistical analyses

Values are represented as means and standard deviations. All statistical tests were performed with GraphPad Prism software, version 7.0c. Specific statistical tests and replicates are indicated in figure legends. Values were considered statistically significant when *p*-values were \**p* < 0.05; \*\**p* < 0.01, \*\*\**p* < 0.001; \*\*\*\**p* < 0.0001.

## Results

### SL7207 inhibits tumour growth and increases survival

The B16F10 allograft melanoma model is a well-characterised *in vivo* tumour model (Overwijk & Restifo, 2001). To validate the tumour-growth inhibitory effects of SL7207 in this model, tumour-bearing mice were treated with 5 x 10^6^ CFU of SL7207 in cold PBS or PBS alone via intravenous injection and tumour growth was measured for 7 days. SL7207 significantly inhibited tumour growth compared to the PBS control, and indeed the size of the tumour 7 days post SL7207 administration was not significantly increased compared to the day of infection (Figure 1A). Furthermore, there was a significantly lower fold growth of the infected tumours compared to the uninfected PBS controls from the time of treatment to the time of harvest (Figure 1B). There was also a significant improvement in survival of the infected mice compared to PBS treated controls at 14 dpi (30 days after tumour induction; Figure 1C), however this was also accompanied by weight loss in the infected group (Figure 1D). Finally, CFU counts of SL7207 recovered from various organs demonstrated that the total number of bacteria in the tumours increased from 1-day post infection (1 dpi) and between 3 and 9 dpi, followed by a decrease at 11 dpi (Figure 1E). There were significantly greater numbers of bacteria recovered from the tumours compared to the liver or spleen at 9 dpi.

**Figure 1.**
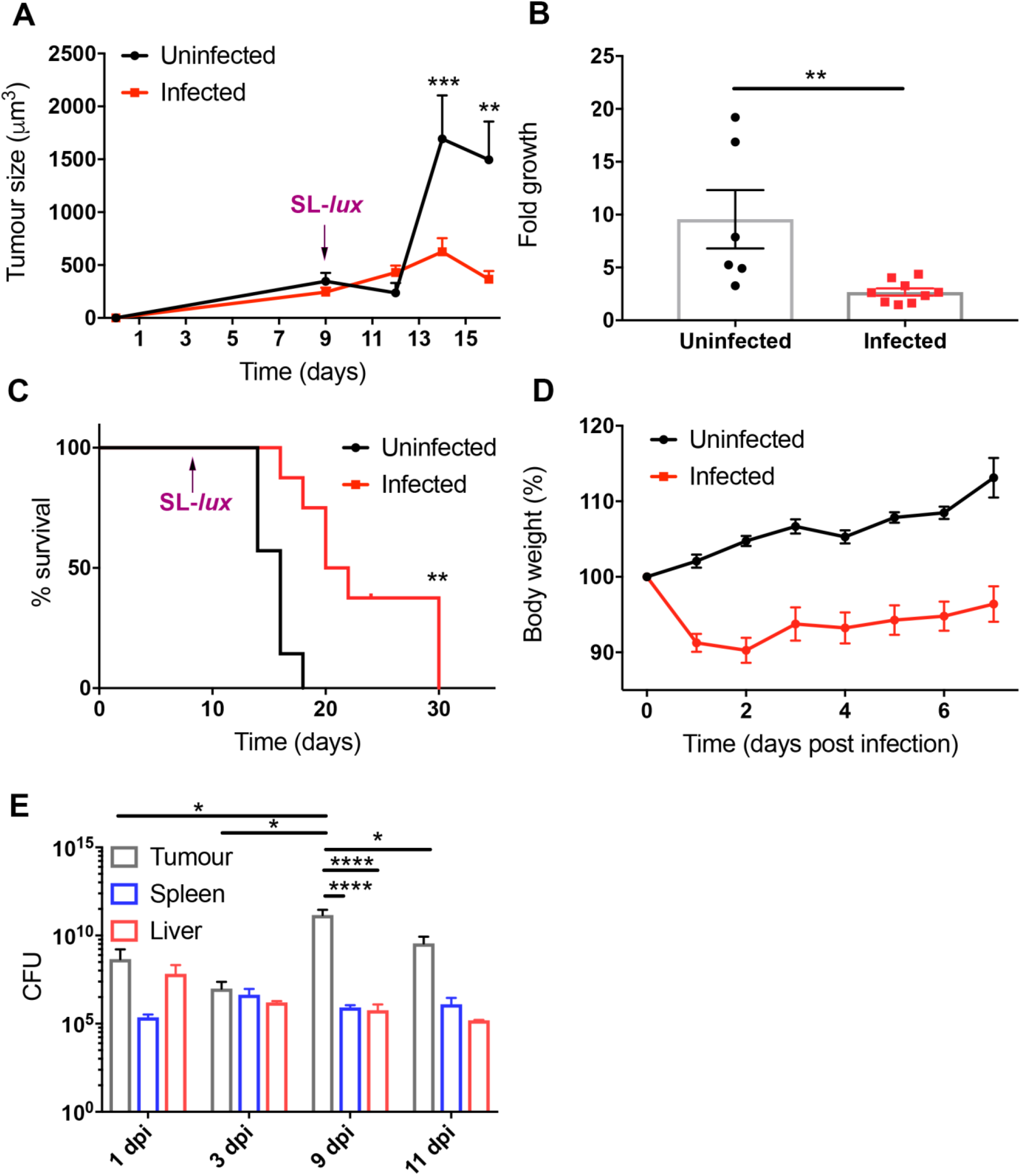
SL7207 inhibits tumour growth in B16F10 tumour models. (**A**) B16F10 tumours were allowed to develop in C57BL/6 mice. Serial tumour size measurements were taken with Vernier calipers at indicated time points (n= 4). Tumour size was calculated using 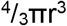 adapted from (Crull *et al*., 2011). (**B**) Fold growth of tumours at the time of harvest 5 dpi compared to tumour size at the time of infection (n= at least 6). (**C**) Kaplein Meier survival curve of tumour-bearing mice infected with SL7207 versus uninfected (purple arrows indicate time point of SL-*lux* administration) (n= 6; endpoint = 14 dpi). (**D**) Weight of mice expressed as a percentage of weight at Day 0 of infection (n= 6). (**E**) Total CFUs of SL7207 at multiple time points in tumours, livers and spleens of infected mice (n= 4). Results displayed are from at least two independent experimental replicates. Results are displayed as the mean ± SD with each point representing a single animal. Samples were analysed using a two-way ANOVA with Sidak post-test correction (**A**), Student’s t-test (**B**), Log Rank Mantel-Cox test (**C**) or using a two-way ANOVA with Tukey post-test correction, (**D** and **E**) where \**p*<0.05; \*\**p*<0.01; and \*\*\**p*<0.001.

### SL7207 administration induces a pro-inflammatory response in the tumour

In order to assess the immune profile of the tumour following systemic infection of SL7207, tumour-bearing mice were infected with SL7207 or control PBS and tumours were harvested and subjected to flow cytometry or ELISA analysis. We found that there was a significant increase in the density of CD45^+^ immune cells at both 5 and 7 dpi (Figure 2A and B; see gating strategy in Supplementary Figure S1). Furthermore, there was a significant increase in the amount of pro-inflammatory cytokines IFN-γ and TNF-α at 5 dpi in the infected group compared to uninfected, with TNF -α maintaining this significant increase at 7 dpi (Figure 2C and D).

**Figure 2.**
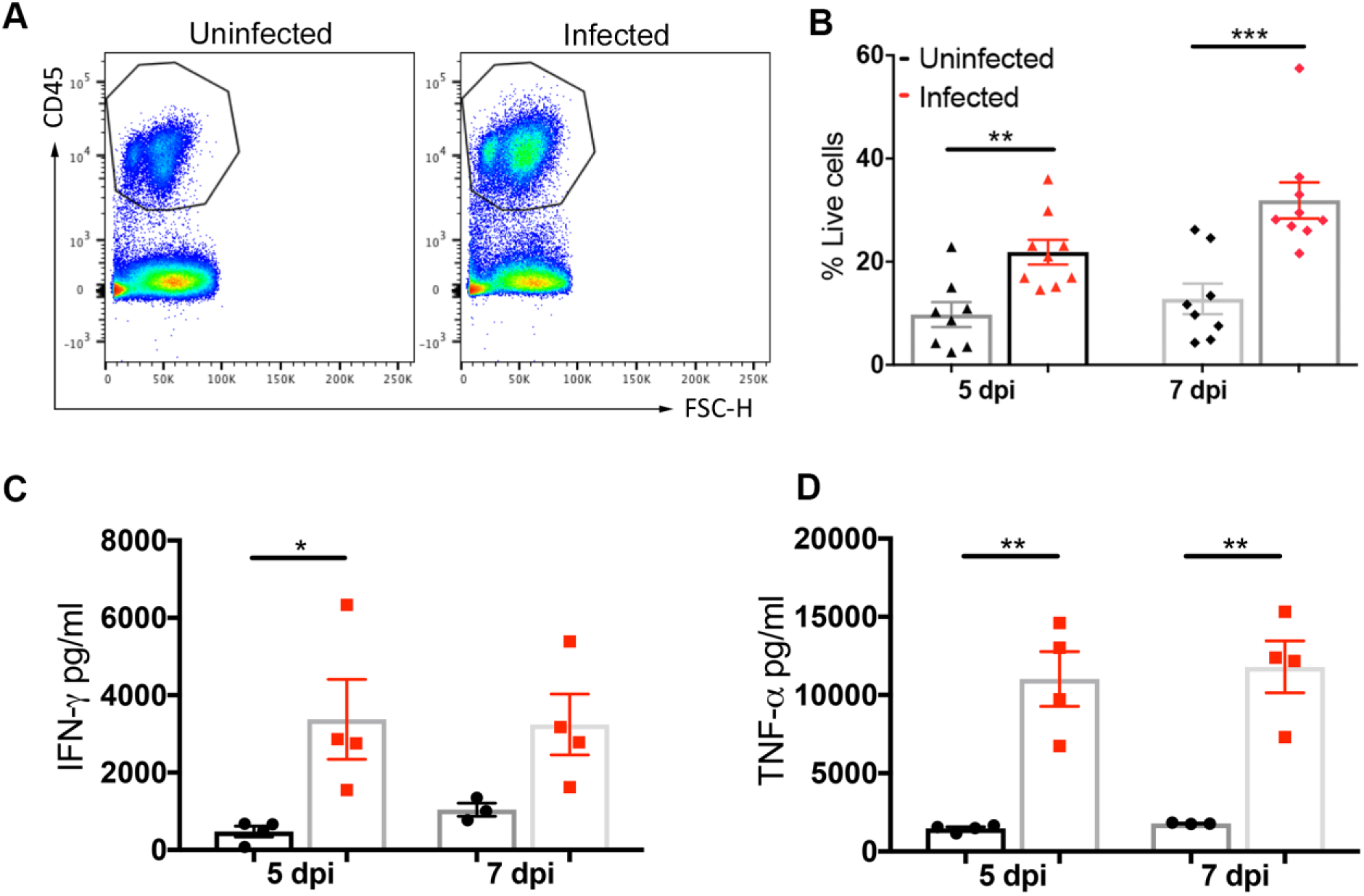
Systemic administration of SL7207 induces immune cell infiltration and pro-inflammatory immune response in the tumour. (**A**) Representative flow cytometry plots of CD45^+^ cells in tumours. Cells were gated on live, single cells. (**B**) Percentage of live cells that are CD45^+^ (n= at least 8). (**C**) ELISA analysis of tumour lysates for IFN-γ and (**D**) TNF-α (n= 4). Results displayed (**A, C** & **D**) are representative of two independent experiments; results displayed (**B**) are from two independent experimental replicates. Results are displayed as the mean ± SD with each point representing a single animal. Samples were analysed using a Student’s t-test where \**p*<0.05; \*\**p*<0.01 and \*\*\**p*<0.001.

### SL7207 infection is accompanied by an influx of pro-inflammatory monocytes into the tumour

Given the significant increase in the inflammatory status of the tumour following infection, it was pertinent to identify which immune cell types were contributing to this phenomenon. Therefore, tumours from mice infected with SL7207 or administered with PBS for 7 days were subjected to flow cytometry analysis for specific immune cell populations. Of particular interest was the change to the Ly6C^+^ monocyte compartment (Figure 3A). More specifically, although there was no change in the proportion of Ly6C^+^MHCII^-^ monocytes, these cells exhibited increased proliferation (Ki67^+^) and secretion of multiple pro-inflammatory cytokines (Figure 3B). Furthermore, Ly6C^+^MHCII^+^ monocytes exhibited a significant expansion within the CD45^+^ immune cell compartment following infection, and these cells exhibited increased proliferation and cytokine secretion compared to PBS-treated controls (Figure 3C). The proportion of Ly6C^-^MHCII^+^ macrophages remained unchanged between infected and uninfected samples, as did their production of pro-inflammatory cytokines, however there was an increase in the proliferative capacity of these cells (Supplementary Figure S2). Finally, there appeared to be a decrease in the proportion of Ly6C^-^ MHCII^-^ macrophages in the CD45^+^ population for the infected samples compared to uninfected samples (Supplementary Figure S2). However, there were no differences in the production of pro-inflammatory cytokines between the two groups of mature macrophages, although the infected samples exhibited a greater proliferative capacity compared to uninfected controls (Supplementary Figure S2).

**Figure 3.**
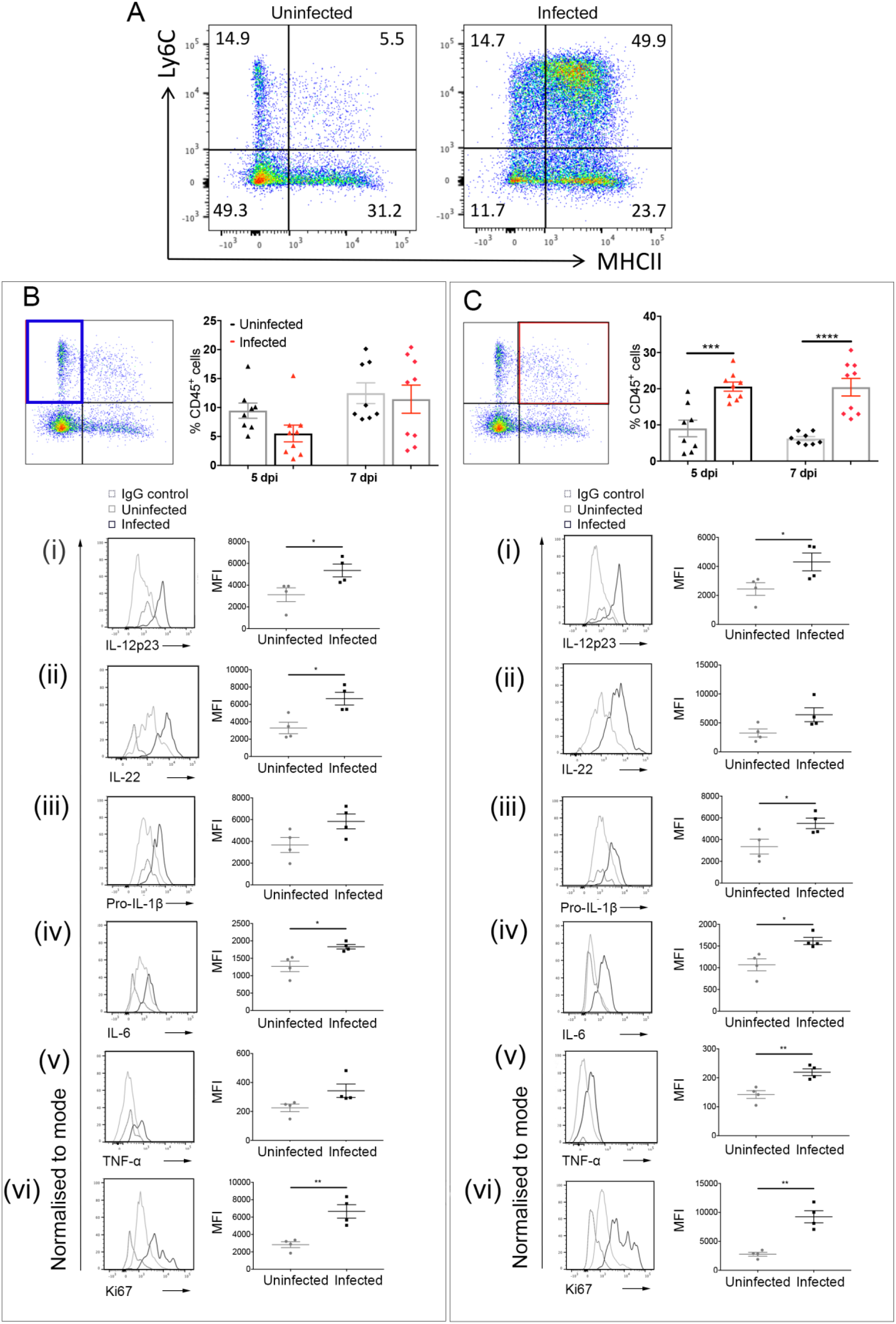
Administration of SL7207 induces Ly6C^+^MHCII^+^ monocyte accumulation accompanied by increased expression of pro-inflammatory markers. (**A**) Representative flow cytometry plots of monocyte and macrophage populations at 7 dpi. Cells gated on live, single cells, CD45^**+**^, CD11b^**+**^, SiglecF^**-**^, Ly6G^**-**^, F4/80^**+**^. (**B**) Depiction of relevant cell population (Ly6C^**+**^MHCII^**-**^ monocytes; left panel) with number of cells as a percentage of CD45^**+**^ cells at 7 dpi (right panel; n= at least 8) and the relative expression of (**i**) IL-12p23, (**ii**) IL-22, (**iii**) pro-IL-1β, (**iv**) IL-6, (**v**) TNF-α and (**vi**) Ki67 in this population with a representative plot from each sample group: isotype control (broken grey line), uninfected (light grey line) and infected (dark grey line). (**C**) Depiction of relevant cell population (Ly6C^**+**^MHCII^**+**^ monocytes; left panel) with number of cells as a percentage of CD45^**+**^ cells at 7 dpi (right panel; n= at least 8) and the relative expression of (**i**) IL-12p23, (**ii**) IL-22, (**iii**) pro-IL-1β, (**iv**) IL-6, (**v**) TNF -α and (**vi**) Ki67 in this population with a representative plot from each sample group: isotype control (broken grey line), uninfected (light grey line) and infected (dark grey line). Representative flow cytometry plots from two mice representative of two independent experiments (**A**, **B** & **C**); all graphs (**B** & **C**) show results from two independent experimental replicates; all plots (**Bi-vi** & **Ci-vi**) show quantitative data from one experiment (n= 4) representative of two independent experiments. Results are displayed as the mean ± SD with each point representing a single animal. Samples were analysed using a Student’s t-test where \**p*<0.05; \*\**p*<0.01; \*\*\**p*<0.001; and \*\*\*\**p*<0.0001.

### Administration of clodronate interferes with the tumour-growth inhibitory effects of SL7207

Given the increase in production of IL-1β, TNFα, IL-12, IL-22, and IL-6 by the expanded inflammatory Ly6C^+^MHCII^+^ monocyte population in the tumour following the administration of SL7207, it was hypothesized that monocyte populations may underpin the tumour-growth inhibition effects of SL7207. Therefore, clodronate liposome treatment, which depletes monocyte populations *in vivo* whilst sparing other phagocytic populations (Supplementary Figure S3), was employed. Tumour-bearing mice were treated with clodronate liposomes or PBS liposomes at multiple time points over the course of the experiment (Figure 4A) in line with previously published protocols (Gazzaniga *et al*., 2007; Griesmann *et al*., 2017). Mice were also either infected with SL7207 or treated with control PBS, and tumour growth was monitored for 7 days before tumours were harvested for analysis. Tumour-bearing mice which were treated with PBS liposomes and subsequently infected with SL7207 showed significantly reduced tumour growth compared to uninfected PBS control mice at 7 dpi (*p* < 0.0001). However, in the clodronate liposome-treated groups, the infected tumours grew similarly to the uninfected, PBS control tumours (*p* = 0.7929). Furthermore, the tumours from the clodronate liposome-treated infected mice were significantly larger in size than the PBS liposome-treated infected tumours (Figure 4A and B; *p* = 0.0001). The clodronate liposome-treated infected mice did not have the same degree of weight loss as the PBS liposome-treated infected mice (Figure 4C). Thus, it appears that treatment with clodronate liposomes abrogated the anti-tumour effect mediated by SL7207 treatment. Finally, to rule out the possibility that the liposome treatment had an effect on bacterial colonisation of the tumours in infected mice, tumours were collected from all groups at 7 dpi and bacterial CFUs were compared. There were no significant differences in the CFU counts between experimental groups in any of the organs at this time point (Figure 4D).

**Figure 4.**
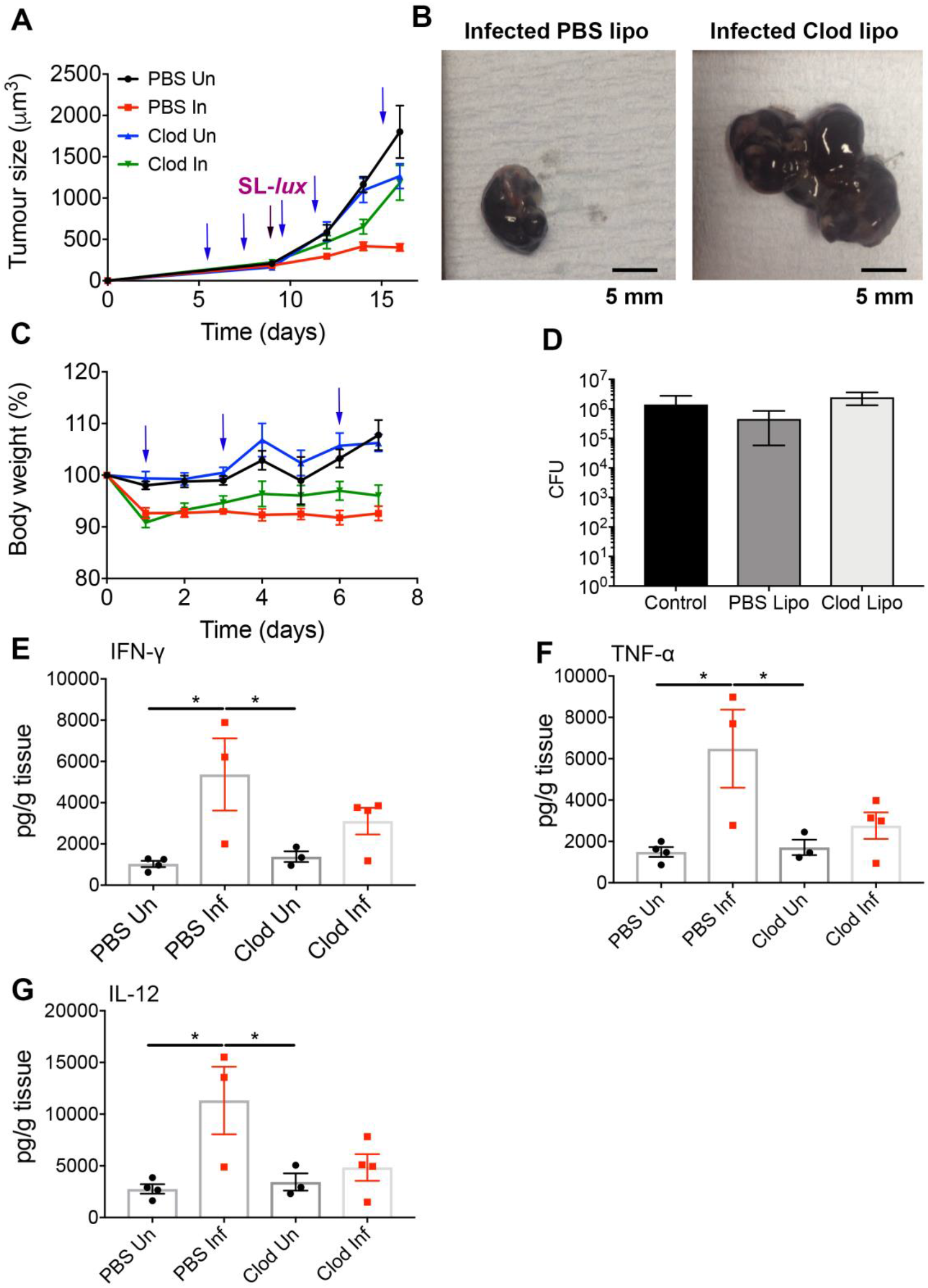
Clodronate liposome (Clod lipo) administration protects tumours from SL7207-mediated growth inhibition. (**A**) B16F10 melanoma tumours were allowed to develop in C57BL/6 mice (n= at least 3) with serial measurements and were subjected to PBS Lipo administration or Clod Lipo administration (blue arrows) with or without SL7207 infection (purple arrow). (**B**) Representative photographs of tumours from infected PBS Lipo and Clod Lipo tumours at 7 dpi (scale bar 5 mm). (**C**) Weight of mice in different groups (refer to Figure 4A) expressed as a percentage of weight at Day 0 of infection/PBS administration following treatment with PBS Lipo or Clod Lipo (blue arrows) (n = at least 3). (**D**) CFU of tumours for indicated conditions at the time of harvest with ‘control’ referring to no liposome treatment (7 dpi) (n= 3). (**E**) ELISA analysis of tumour lysates at 7 dpi for IFN-γ and (**F**) TNF-α. (**G**) ELISA analysis of tumour lysates at 7 dpi for IL-12p23. Results displayed (**A** & **C**) are from two independent experimental replicates; results displayed (**D**) are from one experiment representative of two independent experimental replicates; results displayed (**E-F**) are from one experimental replicate (n= at least 3). Results are displayed as the mean ± SD with each point representing a single animal. Samples were analysed using two-way ANOVA with Sidak post-test correction (**A**), two-way ANOVA with Tukey post-test correction (**C**) or a one-way ANOVA (**D-G**) where \**p*<0.05.

### Treatment with clodronate liposomes decreases the pro-inflammatory status of the tumours following infection

As it was hypothesized that the pro-inflammatory monocytes were mediating the anti-tumour effects of SL7207, it was pertinent to investigate the inflammatory status of the tumour following liposomal treatments with and without infection. This was achieved by harvesting the tumours at 7 dpi and measuring the cytokines, IFN-γ, TNF-α and IL-12 by ELISA. There was a significant increase in the production of all three cytokines by the PBS liposome-treated, infected samples compared to the uninfected controls (Figure 4E-G). However, this increase in cytokine levels in infected samples was hindered for the clodronate liposome and infected samples compared to uninfected controls.

## Discussion

The ability of bacteria, in particular *Salmonella enterica*, to induce tumour growth inhibition is well documented in the literature (Avogadri *et al*., 2008; Crull *et al*., 2011; Kocijancic *et al*., 2017; Lee *et al*., 2011). However, there remain numerous questions as to the mechanisms underlying this effect. Some studies have focused on metabolic factors, others on migratory factors but also many on the immune response mediated following infection. However, in those that examine the immune response, roles have been suggested for neutrophils, dendritic cells and T-cell subtypes in contributing to the tumour growth inhibitory effects of bacteria (Avogadri *et al*., 2008; Kocijancic *et al*., 2017; Stern *et al*., 2015; Westphal *et al*., 2008). Here, we focused on the contribution of the monocyte compartment to the anti-tumour effects of the attenuated *Salmonella enterica* serovar Typhimurium strain SL7207, a strain known to slow tumour growth *in vivo*.

While it has previously been demonstrated that SL7207 has an anti-tumour effect in multiple *in vivo* models (Crull *et al*., 2011; Stern *et al*., 2015), it was pertinent to validate the effects of SL7207 in our model. The melanoma model employed in this study is based on the use of metastatic melanoma cells, thus these tumours tend to grow more quickly than other models, restricting the time frame of the experiments presented herein to approximately 16 days from the initial tumour cell administration to the mice. Initial work showed that the model was sufficient to investigate the anti-tumour and pro-survival effects of SL7207, as well as weight loss and infection in other organs (Figure 1).

The tumour microenvironment is one of an immunosuppressive phenotype, which limits the ability of the immune system to reject the tumour (Rodriguez *et al*., 2004). Tumours are often host to a high density of regulatory T cells (T_reg_) which arrest anti-tumour immune responses using inhibitory receptors such as PD-1 and CTLA-4, resulting in the inhibition of antigen presenting cells (Strauss *et al*., 2007; Uhlig *et al*., 2006). Furthermore, these cells and others produce immunomodulatory molecules such as IL-10, IL-4 and TGF-β (Chen *et al*., 2005; Gorelik *et al*., 2002; Murai *et al*., 2009). In the present study, it is demonstrated that the administration of SL7207 is accompanied by an influx of immune cells into the tumour environment. Furthermore, these immune cells contribute cytokines such as TNF-α (Yrlid *et al*., 2000), which is known to have anti-tumour effects *in vivo* (Sabel *et al*., 2004). These findings are consistent with human studies. In one of these, in patients with colorectal cancer, patients with high T_H_1 cytokine production, such as IFN-γ and TNF-α, had prolonged survival (Tosolini *et al*., 2011).

It has been demonstrated that in up to 80% of clinical studies of cancer, increased macrophage density correlated with poor prognosis (Bingle *et al*., 2002). Tumour-associated macrophages are implicated in tumour progression in multiple ways. Tumour-associated macrophages can produce immunomodulatory cytokines (Facciabene *et al*., 2011; Saccani *et al*., 2006), express immunoinhibitory molecules (Belai *et al*., 2017; Gordon *et al*., 2017; Noman *et al*., 2014), stimulate tumour neovascularization (Coffelt *et al*., 2010; De Palma *et al*., 2005) and suppress T cell function (Bak *et al*., 2008; Rodriguez *et al*., 2004). It has been shown that tumour-associated macrophages derive from circulating CCR2^+^ bone-derived monocytes (Franklin *et al*., 2014). Both monocytes and macrophages are activated following inflammation and bacterial infection (Bain *et al*., 2013; Yrlid *et al*., 2000)). The acquisition of Ly6C^hi^ monocytes is increased following inflammation in the colon (Bain *et al*., 2013; Zigmond *et al*., 2012) and these cells, which normally give rise to tissue macrophages, halt in their differentiation pattern and remain as Ly6C^+^MHCII^+^ “intermediate” cells. Furthermore, these cells appear to have greater TNF-α production than mature macrophages and are more responsive to toll-like receptor stimulation, suggesting that they are prime contributors to inflammation (Bain *et al*., 2013). This phenomenon is also seen in the present study, whereby there is an accumulation of Ly6C^+^MHCII^+^ monocytes in the tumours following infection. These cells appear to be the principal contributors to the production of inflammatory cytokines in the monocyte/macrophage compartment following infection as neither of the tumour-associated macrophage populations, MHCII^-^ nor MHCII^+^, produced greater quantities of pro-inflammatory markers with infection compared to PBS controls. This observation suggests that the mature macrophages in the tumour are not capable of switching their phenotype towards a T_H_1-cytokine producing state as has been suggested in previous publications (Guiducci *et al*., 2005). Taken together, these data suggest that the inflammatory state demonstrated in the tumour following infection is most likely due to the accumulation of monocytes that are recruited following infection. To our knowledge, the present study is the first to describe this phenomenon in tumour inflammation. Other studies on bacterial-mediated cancer treatment have suggested a role for macrophages in mediating anti-tumour effects (Lizotte *et al*., 2014; Zheng *et al*., 2017). The data provided here suggests that it was likely the recently recruited monocytes, not more mature macrophages as was previously suggested, which were the primary contributors. In the present study, the requirement of monocytes was investigated with clodronate liposomes, which target phagocytic cells, particularly monocytes and macrophages. The administration of clodronate liposomes abrogated the tumour growth inhibitory effects of SL7207 highlighting the necessity of these cells to mediate the anti-tumour effects of the bacteria. To our knowledge, this is the first study to highlight the necessity of these cells to do so.

These findings are of therapeutic interest for cancer treatment. Firstly, they reiterate the important contribution of monocytes and macrophages to the tumour microenvironment, particularly the impact these cells can have on making this environment anti-tumourigenic when they are suitably stimulated. Secondly, it suggests that these cells may be manipulated to slow tumour growth. This could be achieved by bacteria, as has been shown here, or perhaps also with other monocyte-activating agents. Finally, this study contributes to our understanding of the immune responses required to mediate effective cancer immunotherapy, providing an opportunity for further investigation into these cells in future studies.

## Author Contributions

S.A.J. was awarded the funding, designed and performed the experiments, analyzed the data, and prepared the manuscript. M.J.O. assisted in experimental design, animal experiments, and ELISA. HMW assisted in experimental design and animal experiments. HEH and ABB assisted with animal experiments. AM provided technical assistance throughout. SM assisted with animal experiments and provided training. SBC, SWGT, AMM and SWFM assisted with experimental design and data analysis and provided reagents. K.B. was awarded the funding, assisted in experimental design, analyzed the data and provided reagents. D.M.W. was awarded the funding, developed the initial concept, designed experiments, and prepared the manuscript. All authors contributed to editing the manuscript for publication.

## Funding

This work was funded by the Wellcome Trust through a Wellcome Trust PhD studentship to SAJ (102460/Z/13/Z) and Biotechnology and Biological Sciences Research Council grants (BB/K008005/1 & BB/P003281/1) and a Tenovus Scotland grant to DMW; and Cancer Research UK grants (CRUK A17196 and A29799) to KB.

## Acknowledgements

We would also like to thank staff at the University of Glasgow Central Research Facility for animal husbandry and at the Institute of Infection, Immunity and Inflammation’s Flow Cytometry Core Facility. We would like to acknowledge the Biological Services Unit at the Cancer Research UK Beatson Institute (C596/A17196).

## Supplementary Figures

**Figure S1.**
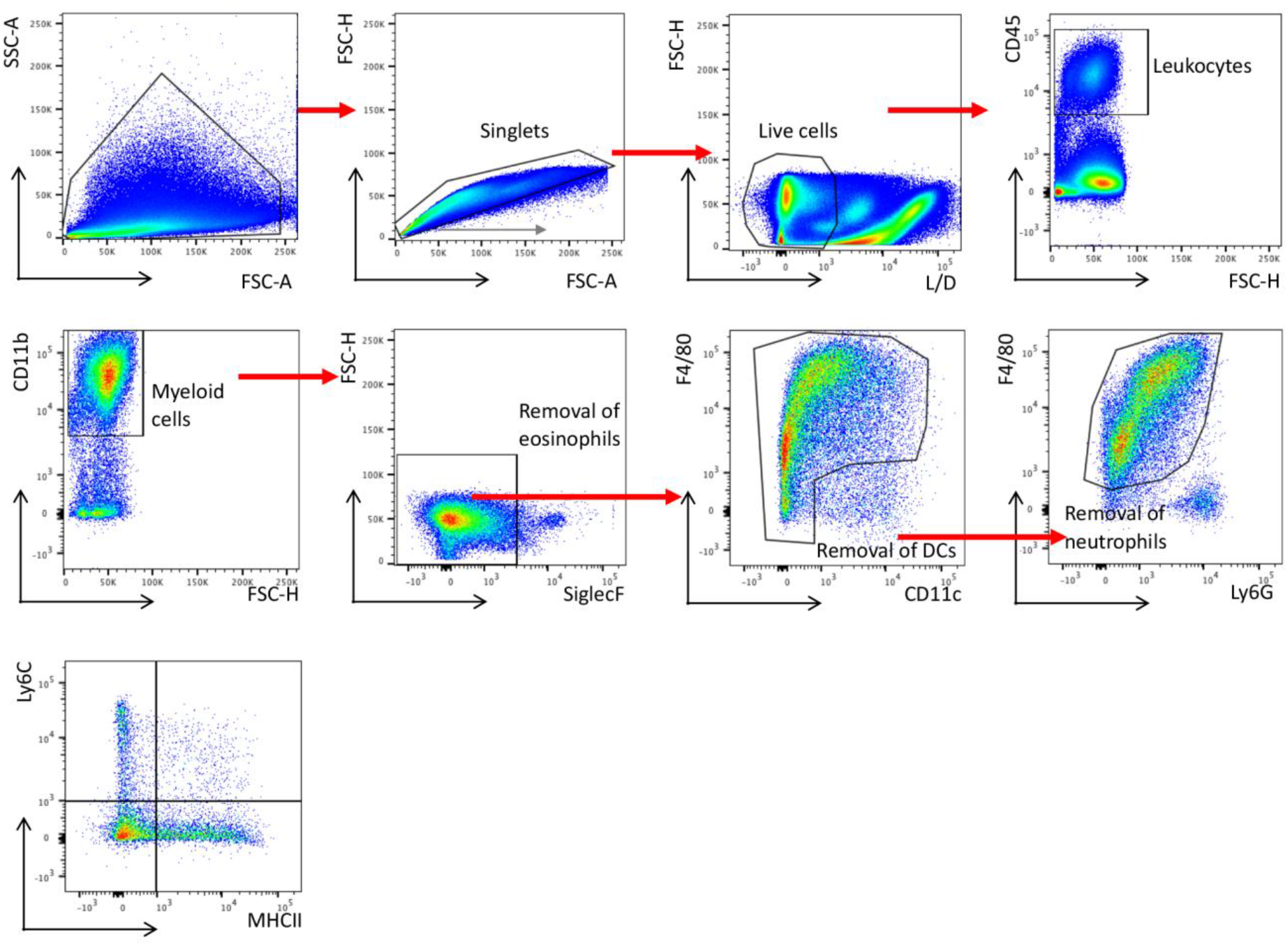
Gating strategy for the monocyte/macrophage compartment. Cells were gated as single (side scatter (SSC) versus forward scatter (FSC)), live, CD45^+^, CD11b^+^, SiglecF^-^ (to exclude eosinophils), CD11c^-^ (to remove dendritic cells), F4/80^+^, Ly6G^-^ (to exclude neutrophils) and thereafter designated according to expression of Ly6C^+^ (monocytes) or Ly6C^-^ (macrophages) and on MHCII expression.

**Figure S2.**
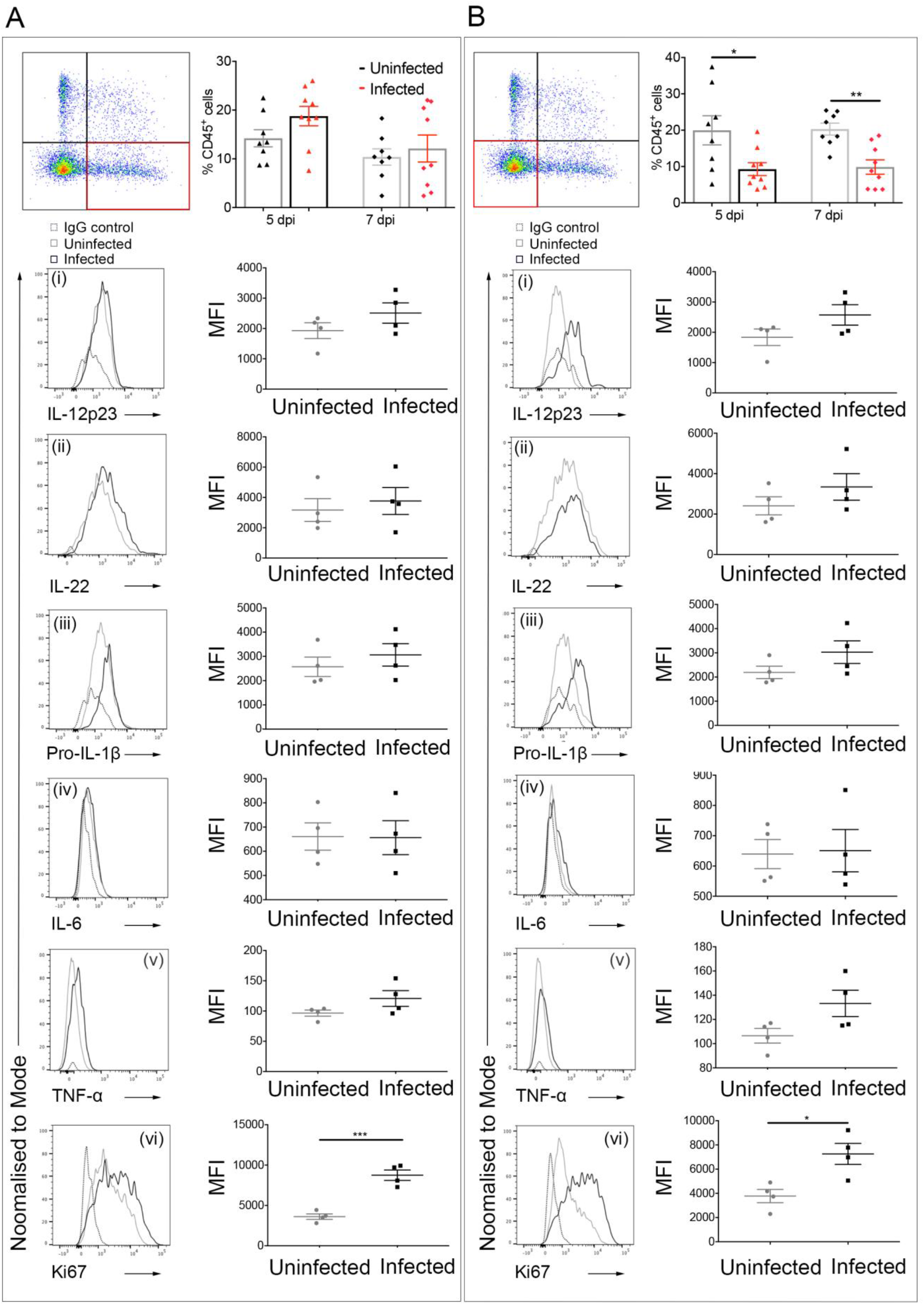
Changes in inflammatory profile of mature TAMs following infection. These data are related to Fig. 3. (**A**) Depiction of relevant cell population (Ly6C^-^MHCII^+^ macrophages; left panel) with number of cells as a percentage of CD45^+^ cells (right panel; n= at least 8) and the relative expression of (**i**) IL-12p23, (**ii**) IL-22, (**iii**) pro-IL-1β, (**iv**) IL-6, (**v**) TNF-α and (**vi**) Ki67 in this population with a representative plot from each sample group: isotype control (broken grey line), uninfected (light grey line) and infected (dark grey line). (**B**) Depiction of relevant cell population (Ly6C^-^MHCII^-^ macrophages; left panel) with number of cells as a percentage of CD45^+^ cells (right panel; n= at least 8) and the relative expression of (**i**) IL-12p23, (**ii**) IL-22, (**iii**) pro-IL-1β, (**iv**) IL-6, (**v**) TNF -α and (**vi**) Ki67 in this population with a representative plot from each sample group: isotype control (broken grey line), uninfected (light grey line) and infected (dark grey line). Cell plots displayed (**A** & **B**) show data representative of two independent experiments; all graphs (**A** & **B**) show results from two independent experimental replicates ± SD; all plots (**Ai-vi** & **Bi-vi**) show quantitative data from one experiment representative of two independent experiments. Samples were analysed using a Student’s t-test where \**p*<0.05; \*\**p*<0.01; \*\*\**p*<0.001; and \*\*\*\**p*<0.0001.

**Figure S3.**
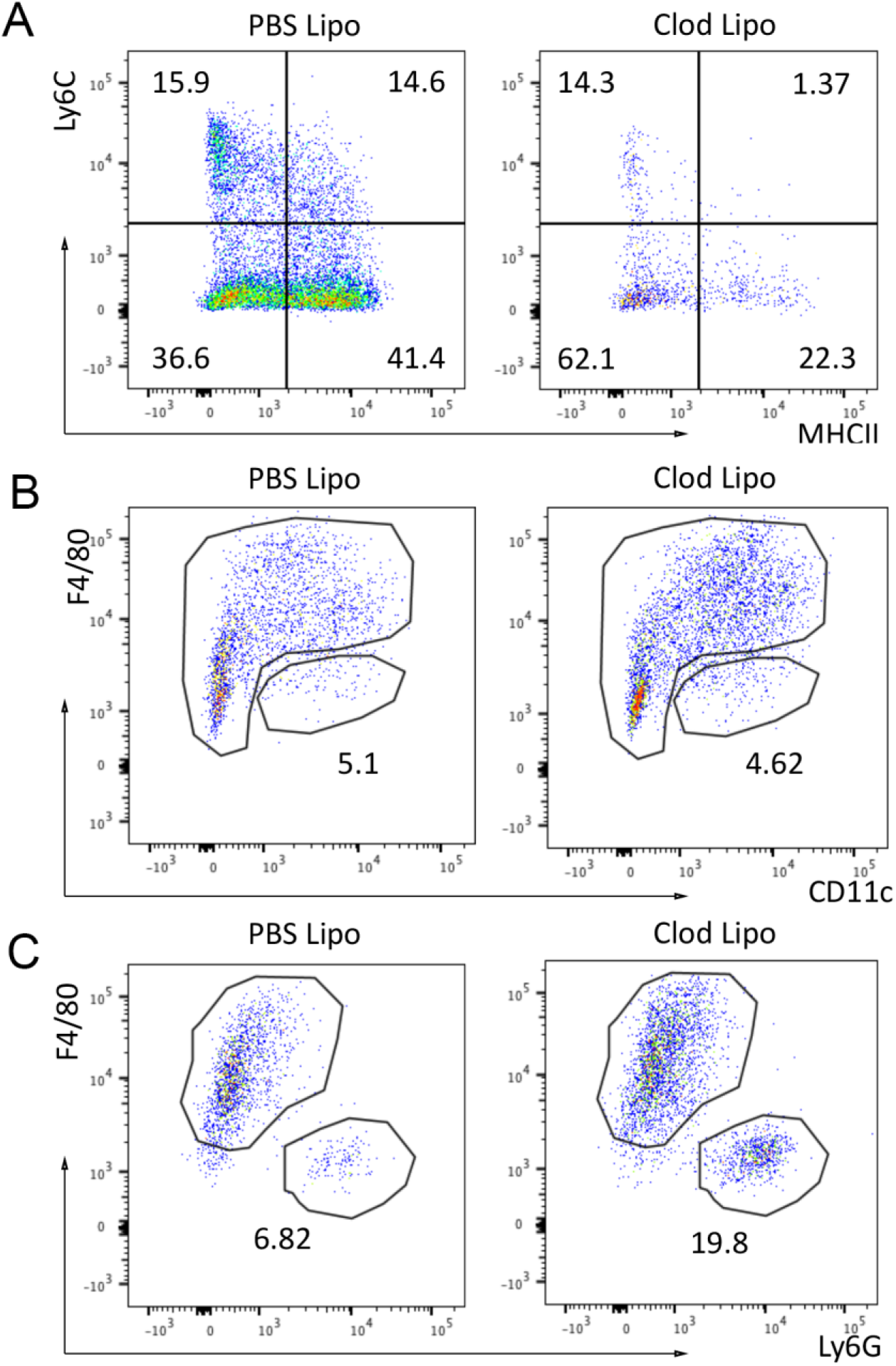
Effects of clodronate liposomes on immune cell populations. (**A**) Flow cytometry analysis of the monocyte/macrophage compartment in tumours following administration of clodronate liposomes or PBS liposomes. (**B**) Flow cytometry analysis of the neutrophils in tumours following administration of clodronate liposomes. (**C**) Flow cytometry analysis of dendritic cells in tumours following administration of clodronate liposomes. Results displayed (**A**, **B** & **C**) are representative of two independent experiments.

## Notes

### Competing Interest Statement

The authors have declared no competing interest.

